# Cryopreservation of *Hydractinia symbiolongicarpus* sperm to support community-based repository development for preservation of genetic resources

**DOI:** 10.1101/2020.11.03.366344

**Authors:** Aidan L. Huene, Matthew L. Nicotra, Virginia M. Weis, Terrence R. Tiersch

**Affiliations:** Department of Surgery, Thomas E. Starzl Transplantation Institute, University of Pittsburgh, Pittsburgh, PA, USA; Pittsburgh Center for Evolutionary Biology and Medicine, University of Pittsburgh, Pittsburgh, PA, USA; Department of Immunology, University of Pittsburgh, Pittsburgh, PA, USA; Department of Integrative Biology, Oregon State University, Corvallis, OR, USA; Aquatic Germplasm and Genetic Resources Center, School of Renewable Natural Resources, Louisiana State University Agricultural Center, Baton Rouge, LA, USA

**Author notes:** Corresponding authors (TT), (MN).

## Abstract

*Hydractinia symbiolongicarpus* is an emerging model organism in which cutting-edge genomic tools and resources are being developed for use in a growing number of research fields. However, one limitation of this model system is the lack of long-term storage for genetic resources. Our goal in this study was to establish a generalizable approach to sperm cryopreservation that would support future repository development and could be applied to many species according to available resources. Our approach was to: 1) Assess sperm characteristics and standardize collection and processing; 2) Assess acute toxicity to cryoprotectants, and 3) Evaluate and refine freezing conditions to permit post-thaw fertilization and produce viable offspring. By following this approach, we found that *Hydractinia* sperm incubated in 5% DMSO, equilibrated at 4°C for 20 min, and cooled at a rate of 20°C/min to - 80°C at a cell concentration of 10^8^-10^9^/mL in 0.25-mL aliquots were able to fertilize 150-300 eggs which yielded offspring that could metamorphose into juvenile polyps. In addition, improvements were made for processing sperm using a customized 3-D printed collection system. Other opportunities for improvement include optimizing the volumetric sperm-to-egg ratio for fertilization. Establishing repository capabilities for the *Hydractinia* research community will be essential for future development, maintenance, protection, and distribution of genetic resources. More broadly, this application-based approach highlights the long-term value of establishing repository-level resources that can be expanded to fit community needs.

## Introduction

*Hydractinia symbiolongicarpus* is a colonial cnidarian and an established model for evolutionary developmental biology, stem cell biology, regeneration, and allorecognition [1–3]. In recent years, efforts to improve *Hydractinia* as a model system have included generation of robust laboratory strains for use by the research community, sequencing of these strains through the *Hydractinia* Genome Project (https://research.nhgri.nih.gov/hydractinia/), and establishment of methods to produce transgenic animals via the random integration of exogenous DNA [4] or targeted integration via CRISPR/Cas9-mediated gene knock-in [5].

An increasing limitation to the expanded use of *Hydractinia* as a model is the lack of long-term storage options for genetic resources. Over the years, laboratories have collected and bred hundreds of genotypically distinct colonies, while simultaneously generating strains bearing various transgenes. In all cases, these animals have had to be maintained as live animals or they would be lost. While this is possible because *Hydractinia* colonies can be maintained for decades under laboratory conditions, it is increasingly costly in terms of labor and space. These costs are often minimized by reducing colonies to the smallest possible size, and only expanding them via clonal reproduction when needed for experiments. However, these colonies remain vulnerable to accidents, disease, and improper handling which can result in death and permanent loss of genotypes important to previous and future research.

To address this limitation, we sought to evaluate the feasibility and potential utility of cryopreservation as an archival storage method. As an immediate benefit, cryopreservation would allow “backing-up” animals that are valuable genetic resources. And, as a long-term benefit beyond laboratory use, cryopreserved stocks would allow user groups from across the research community to store and access samples on demand rather than requiring time and resources to grow or collect new animals. While the ultimate goal would be cryopreservation of germplasm and somatic tissues from all life stages, here we focused on *Hydractinia* sperm as the most amenable to cryopreservation based on previous success in corals [6], and the anemone *Nematostella* (Matt Gibson and Shane Merryman, personal communication).

Although much is known about *Hydractinia* embryonic development and the differentiation of *Hydractinia* germ cells [7–9], much less is known about *Hydractinia* germplasm after its release, beyond what is necessary for routine breeding. It is well established that *Hydractinia* are dioecious and have gonozooids (reproductive polyps) that bear multiple gonophores (gamete-filled structures) that release either sperm or eggs. Healthy *Hydractinia* release gametes daily. Researchers typically allow male and female colonies to spawn together in the same water or they collect eggs and sperm separately, then mix them within 30 minutes. Anecdotal evidence suggests waiting longer than 30 minutes decreases the quantity and quality of embryos.

After fertilization, each embryo develops into planula larva (1-4 d) before permanently attaching to the surface and metamorphosizing into a juvenile primary polyp. The animal then grows by extending structures called stolons across the surface, from which additional polyps are produced to create a colony. Colonies become sexually mature within 1-2 months. Under laboratory conditions the number of offspring that are male or female is consistent with a 1:1 sex ratio.

Successful cryopreservation of sperm cells requires the balance of multiple parameters [10]. These include the storage temperature and the time that elapses the time and storage temperature that elapses between sperm collection and freezing, sperm concentration at the time of freezing, choice and concentration of cryoprotectant, cooling method and rate, thawing method and rate, and the conditions under which thawed sperm will be used for fertilization [11]. Here we detail a systematic three-part approach to: 1) determine basic characteristics of *Hydractinia* sperm and standardize collection and processing: 2) test the toxicity of commonly used cryoprotectants, and 3) identify conditions that maximized the likelihood of cryopreserved sperm samples being capable of fertilization after thawing.

## Materials and methods

### Ethics

Animal care is overseen by separate Institutional Animal Care and Use Committees at the University of Pittsburgh and Louisiana State University. *Hydractinia symbiolongicarpus* is a marine invertebrate lacking a central nervous system and is not regulated by specialized guidelines. All animals used in this study were maintained in continuous culture as detailed below.

### Animal care and breeding

Experimental work was performed from February to April, 2019, at the Aquatic Germplasm & Genetic Resources Center (AGGRC) in Baton Rouge with animals transported in 50-mL tubes by overnight shipping from University of Pittsburgh. Colonies were maintained and grown as previously described [5] and cultured for at least 2 weeks before use in experiments. Briefly, colonies were established on 25 mm × 75 mm glass microscope slides and cultured in 38-L (10-gal) aquaria using artificial seawater (ASW) (Instant Ocean Reef Crystals, Spectrum Brands, Blacksburg, VA) at between 29 and 31 ppt, held at 22-23°C, and maintained on an 8h:16h (light:dark) photoperiod. Adult colonies were fed 4-day-old *Artemia* nauplii on Monday, Wednesday, and Friday. On Tuesday and Thursday, colonies were fed a suspension of pureed oysters (fresh caught, shucked, pureed, aliquoted, flash frozen in liquid nitrogen, and stored at -20°C).

In this study, we performed crosses between two half siblings, a male (colony 291-10) and a female (colony 295-8) (Fig S1). Following first exposure to light, male and female colonies were moved into separate bins filled with ASW and placed under supplemental lighting. Gametes released approximately 1 hr after light exposure. Sperm were released in “clouds” or “streams” from individual gonophores (**Error! Reference source not found**.A-C) and were collected and pooled using a Pasteur pipette (**Error! Reference source not found**.D).

Eggs were collected by straining the water from the female bin with a 20-μm cell strainer. For routine breeding and to serve as a positive control for fertilization, 20-30 clouds of sperm were collected from 10 male slides, transferred to a 50-mL conical tube, and brought to a final volume of 15 mL with filtered sea water (FSW, artificial seawater filtered through 0.45 μm Polyethersulfone (PES) membrane Rapid-Flow Sterile Disposable Bottle Top Filters, Thermo Scientific Nalgene, catalog #295-4545). To this were added 400-600 eggs harvested from 8-9 female slides. The final volume was brought to 30 mL with FSW and transferred to a 100-mm polystyrene Petri dish. Within 1 hr, embryos began to cleave and developed into planulae by the following day (**Error! Reference source not found**.E). On day 4 after fertilization, larvae (Fig 1F) were settled by exposure to 100 mM Cesium Chloride (CsCl diluted in FSW) for 4-5 hr until ready for settlement, and were pipetted onto microscope slides and kept in the dark for 1-2 d or until attachment and primary polyps formed.

**Fig 1.**
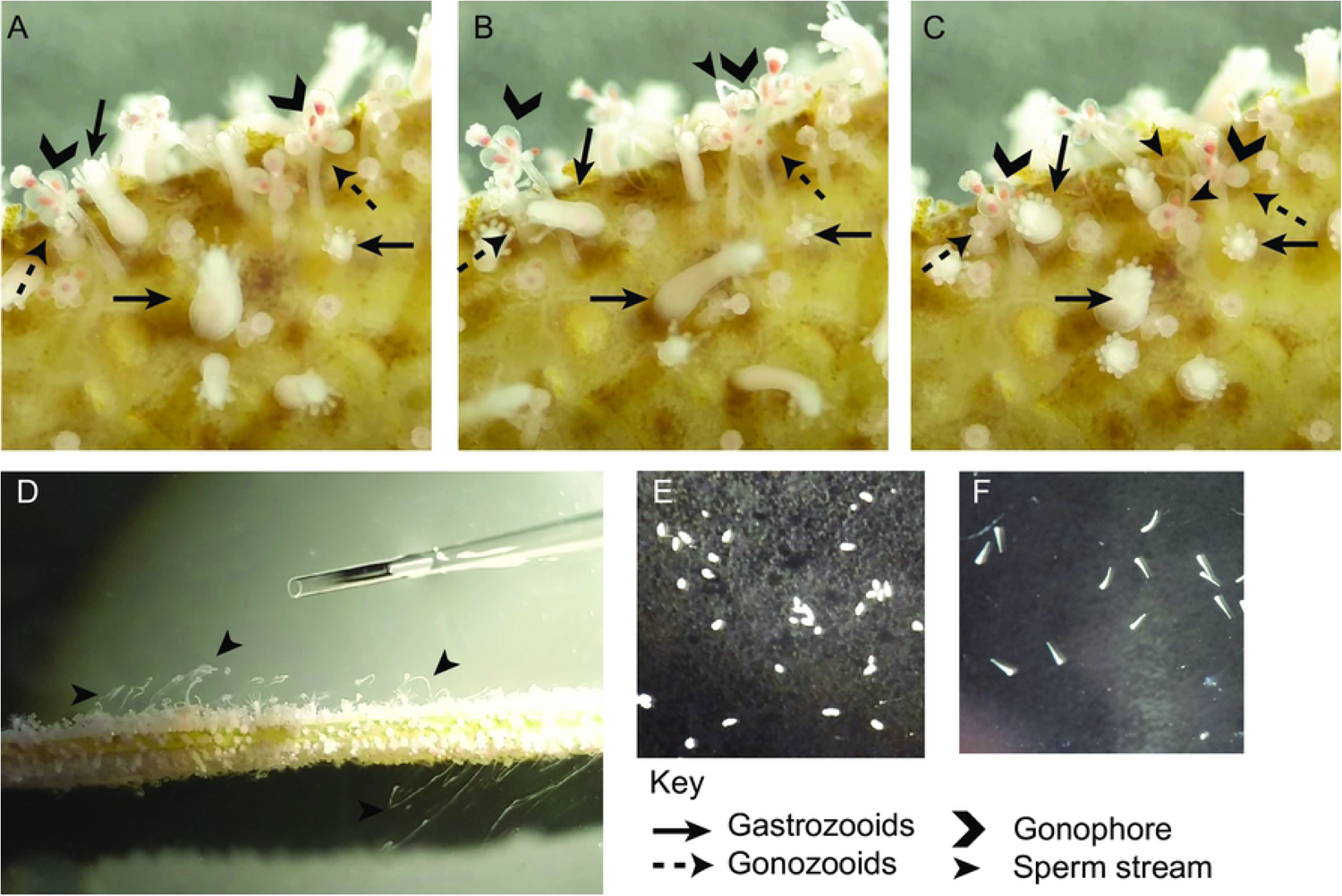
Time lapse of sperm release. (A) Close-up view of Hydractinia polyps just prior to sperm release. Arrows indicate polyp types. (B) Arrowhead points to sperm stream being released. (C) Arrowheads point to sperm stream (polyps have retracted from B). (D) Top-down view of slide with Hydractinia releasing sperm. Arrowheads point to multiple streams of sperm released from the colony. (E) 1-d old larvae. (F) 2-d old larvae.

### Estimation of sperm concentration and motility

On six separate days, individual sperm clouds (cumulative N = 35) were collected in a 10-µl volume and analyzed for motility within 20 min of collection. The sample was briefly vortexed to form a uniform suspension, loaded onto a Makler^®^ counting chamber (SEFI Medical Instruments Ltd, Irvine Scientific, Santa Ana, CA, USA), and viewed with dark-field illumination at 200-X magnification (Olympus CX41RF, Tokyo, Japan). Sperm were already motile when observed and did not require activation. The sample concentration was counted twice according to an established protocol [12] and the average used as the sperm concentration (at 10^6^/mL). Motility was quantified using a computer-assisted sperm analysis (CASA) system (CEROS model; Hamilton Thorne, Inc., Beverly, MA, USA). The settings used were based on a previous study [13]. Briefly, motility and VCL (curvilinear velocity) were measured for 10 sec. Cell detection was set at a minimum of 25 pixels for contrast and 6 pixels for cell size. In each measurement, 100 frames were captured at a rate of 60 frames/sec. Sperm with an average of >20 μm/s measured path velocity (VAP) were counted by the program as being progressively motile. GraphPad Prism (v8.2.0) was used to calculate correlations between sperm characteristics (velocity, percent motile, and concentration).

### Longevity and temperature sensitivity of sperm

To test the effects of time and temperature on sperm motility, approximately 30 sperm clouds were collected using a 10-μL pipette, pooled and diluted to produce a concentration of 2 × 10^7^ cells/mL, and then divided into two tubes. One tube was kept at room temperature (21-23°C) and the other was kept in a 4°C refrigerator. Each treatment was evaluated hourly for presence or absence of motility for 7 hr.

### Fertility of sperm

To determine how long sperm could produce viable offspring when stored at 4°C, we performed a time-series experiment using a single collection of sperm. Approximately 150 clouds of sperm were collected using a Pasteur pipette and stored in a 50-mL conical tube. Concentration was determined as described above. On day 0, 3 mL of this sample (total of 2 × 10^7^ sperm) were used to fertilize 200 eggs in a total volume of 30 mL FSW. The sperm sample was stored at 4°C. On subsequent days, freshly collected eggs were fertilized with 3 mL (2 × 10^7^ sperm) of sperm in 30 mL FSW. Offspring were followed until they metamorphosed into juvenile polyps.

### Acute Toxicity of Cryoprotectants

Approximately 20 sperm clouds were collected using a 10-μL pipette, pooled, and adjusted to a concentration of 1 × 10^7^ sperm/mL using FSW. Three cryoprotectants, methanol (Fisher Scientific, Waltham, MA) dimethyl sulfoxide (DMSO, Fisher Scientific, Waltham, MA), and glycerol (Sigma-Aldrich, St. Louis, MO) were used. For each cryoprotectant, double strength stocks of 10%, 20%, and 30% (v/v) were created using FSW. The sperm and double-strength cryoprotectant were mixed in equal volumes (100 μL:100 μL) resulting in a final sperm concentration of 5 × 10^6^ sperm/mL and final cryoprotectant concentrations of 5%, 10%, or 15%. Sperm were evaluated at 30 min after addition of cryoprotectant (30 min was chosen as a practical total exposure time required for cryoprotectant equilibration and for packaging and handling of the samples). Presence or absence of motility was used as an estimate for toxicity.

### Standardized Sperm Collection (3-D printing)

Based on the difficulties and inefficiencies experienced during pilot experiments working with *Hydractinia* sperm, we designed a custom sperm collection chamber with integrated slide rack to collect and concentrate sperm for downstream applications (Fig 2) by use of free computer-aided design (CAD) online software (Tinkercad, version 4.7, Autodesk, San Rafael, CA). The design was exported as a stereolithography (STL) file and imported into a 3-D printer slicer software (Simplifiy3D, version 4.0, Cincinnati, OH) to control the printing process (Table S1). Collection chambers were printed in black PLA (ZYLtech Engineering, Spring, TX) filament on a stock Prusa i3 MK3 3-D printer (Prusa Research, Czech Republic) (Table S2).

**Fig 2.**
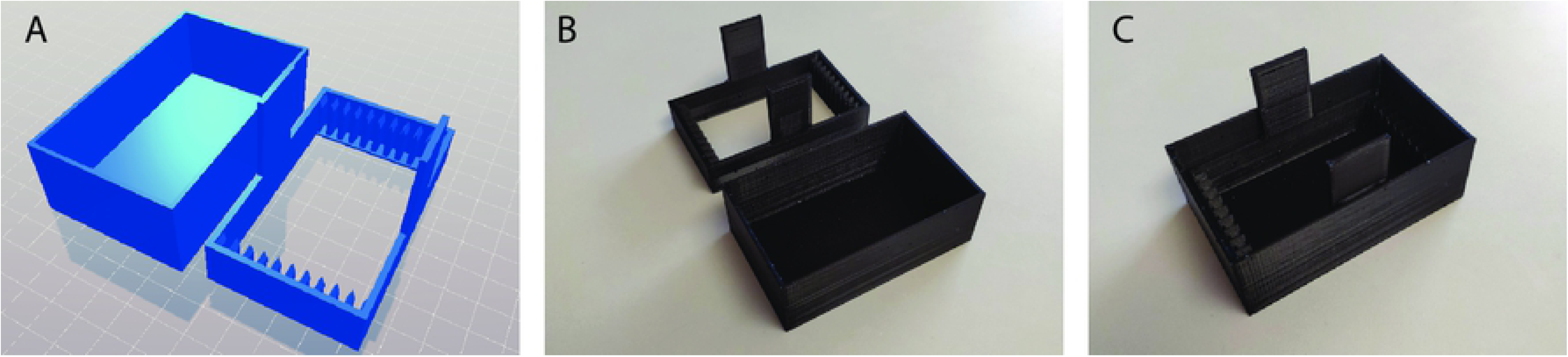
Sperm collection chamber. (A) CAD-rendering of the 3-D design. (B) Printed model with rack and box separate. (C) Printed model with rack inserted. Object model deposited on Thingiverse. https://www.thingiverse.com/thing:3661286

### Freezing

To collect sperm for freezing, we placed nine slides of males in the 3-D printed sperm collection chamber filled with ASW. An additional male was placed in a separate bin so that sperm could be collected and used as a fertilization positive control. After sperm were released, the slide rack was removed from the sperm collection chamber, the cloudy seawater poured into two 50-mL conical tubes (∼80 mL total) and spun for 20 min at 3,000 rpm (∼1450-1500 × *g*) which resulted in a visible white pellet. The supernatant was pipetted off and the pellets were combined and resuspended in FSW to the appropriate concentrations (between 2 × 10^6^ and 2 × 10^9^ sperm/mL) and stored at 4°C until they were prepared for freezing (∼3 hr).

To prepare for freezing, sperm were mixed with an equal volume of 10% DMSO or 10% methanol in FSW (final concentrations of 5% cryoprotectant), drawn into 0.25-mL French straws (IMV International, MN, USA), and held at 4°C in a controlled-rate freezer for the remaining equilibration time (Minitube of America, IceCube 14M, SY-LAB). The total equilibration time, from initial mixing with cryoprotectant to starting the freezing program, was set at 20 min. Equilibrated samples were cooled to -80°C with one of three pre-programmed cooling rates: 5°C/min, 20°C/min, or 30°C/min. Frozen samples were held at -80°C for at least 5 min before transfer and storage in liquid nitrogen.

### Thawing and use for fertilization

After 21-69 hr of storage, straws were removed from liquid nitrogen and immediately plunged into room temperature (22°C) water for 8 sec. The straws were clipped and a 2-μL sample was removed, diluted with 38 μL of FSW (1:20 dilution), and used for sperm assessment. The remaining sample was held in a microfuge tube until fresh eggs were obtained (15-30 min). After performing the fertilization positive control (routine breeding), 100-300 fresh eggs were collected in 500 μL of FSW and added to the microfuge tube with the thawed sperm. The mixed gametes were placed into a 100-mm Petri dish and ∼50 mL FSW was added. An estimate of the number of eggs used was obtained by counting in groups of ten. The resulting fertilization was kept at room temperature and observed for 24 hr to determine how many planulae had begun forming. The resulting offspring were observed until metamorphosis into juvenile polyps.

## Results

### Sperm motility and viability

To assess sperm characteristics, we measured sperm from 35 clouds (each collected in 10 µl) using CASA. Mean velocity was 50.8 ± 26.2 μm/s, mean percent that were motile was 37 ± 22% and mean concentration was 9.37 ± 5.31 × 10^6^/mL. Based on linear regression, velocity and motility were correlated (R^2^ = 0.2804, *P* = 0.0011) (Fig 3A), as were concentration and motility (R^2^ = 0.2870, *P* = 0.0009) (Fig 3B). Velocity and concentration were not correlated (R^2^ = 0.02365, *P* = 0.3778) (Fig 3C).

**Fig 3.**
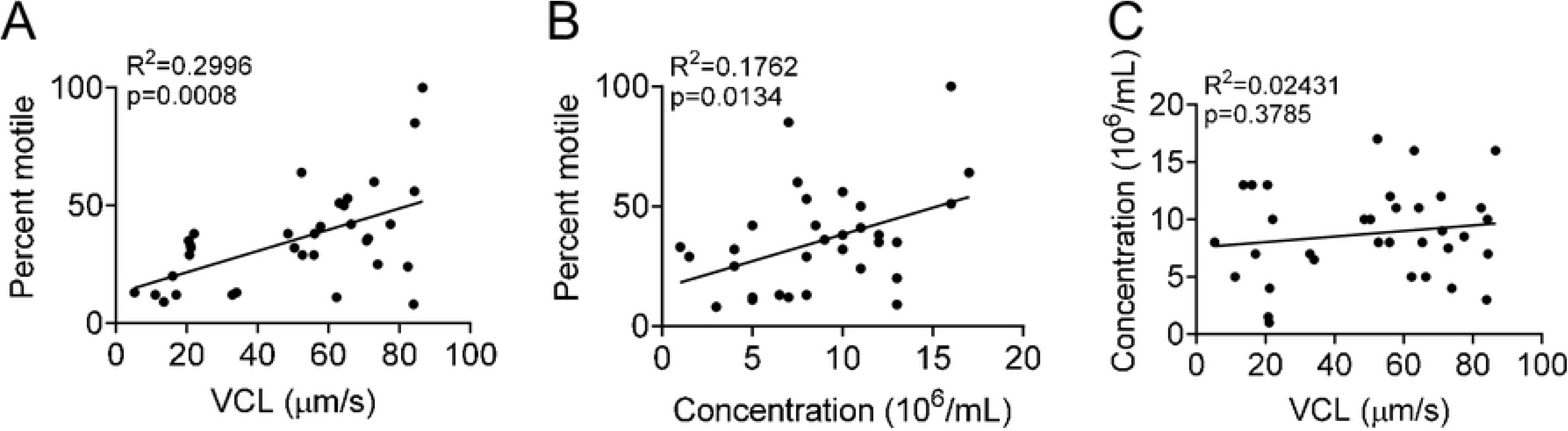
Correlations among sperm velocity, motility, and concentration. Each point represents a sperm cloud (N = 35). (A) Distribution of sperm based on velocity and the number motile. (B) Distribution of sperm based on concentration and number motile. (C) Distribution of sperm comparing velocity and concentration.

To determine the effect of temperature on sperm viability, we compared the motility of freshly collected sperm held at room temperature (22°C) to that of sperm held in a 4°C refrigerator. At room temperature, the number of motile sperm declined over 6 hr, such that by 7 hr only twitching was observed (tail movement without progressive motility). In contrast, sperm kept at 4°C retained progressive motility 7 hr after collection, although the total number of motile sperm and the velocity visibly decreased. By 23 hr, no sperm were motile, but approximately 40%, assessed manually were still twitching. Thus, holding sperm at 4°C prolonged motility.

The observation that sperm held at 4°C were still moving after 23 hr raised the question of whether they could still fertilize eggs and, if so, whether sperm would remain viable after longer storage times. To address this question, we collected ∼150 sperm clouds and used the sperm to fertilize freshly collected eggs over the following 6 d (Fig 4). We performed daily routine breeding to serve as a positive control for egg fertilization; nearly all of the eggs (>95%) were fertilized each day indicating that there was no appreciable differences in egg quality for fertilization. On day 0, we mixed 2 × 10^7^ sperm (3 mL) with ∼200 eggs, which resulted in ∼150 embryos. Because ∼50 eggs remained unfertilized, we interpreted this to indicate that the defined sperm number in this sample (2 × 10^7^) were capable of fertilizing ∼150 eggs.

**Fig 4.**
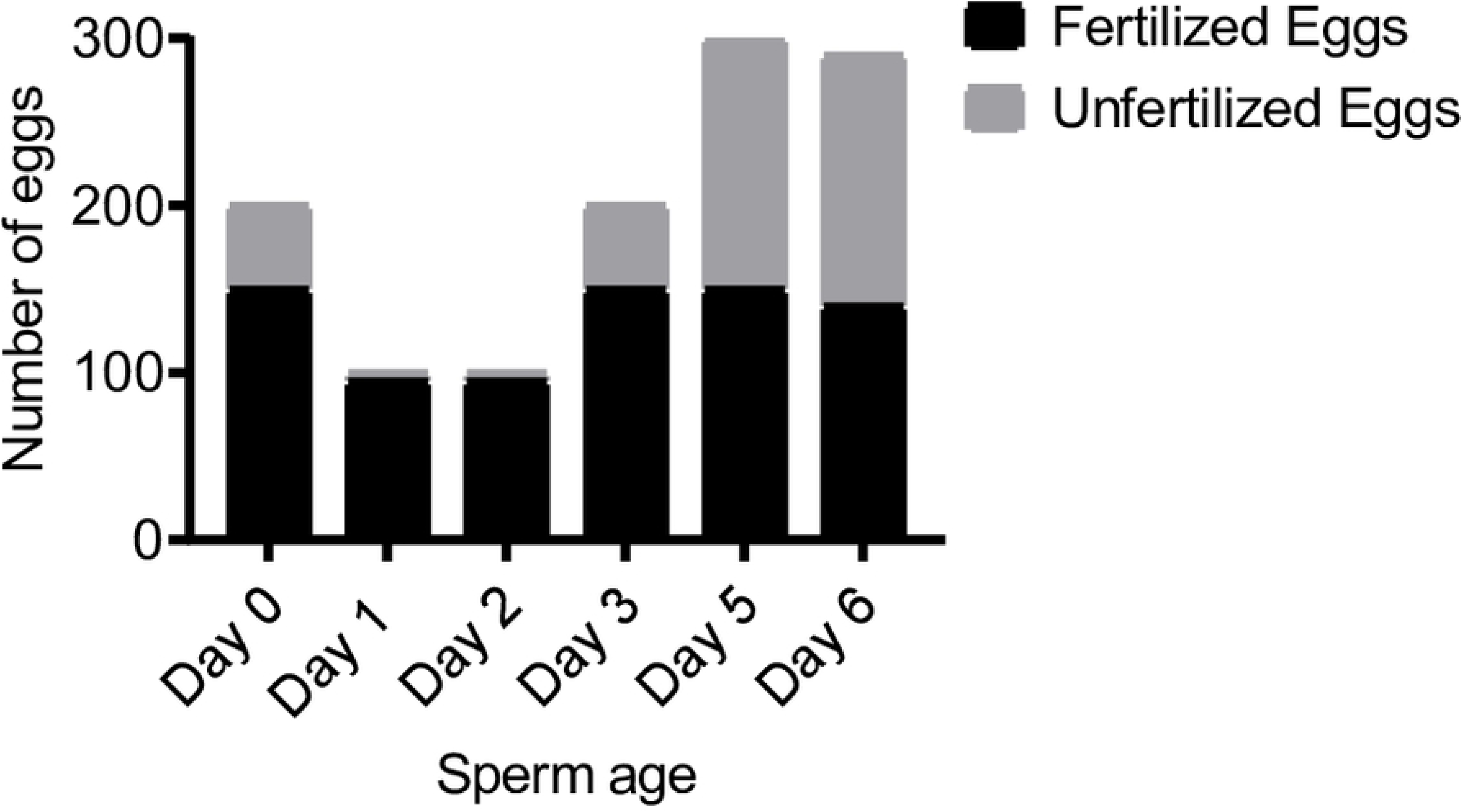
Estimated sperm fertilization capacity over time. Each day, 2 × 10^7^ sperm cells from the same collection aliquot were used to fertilize the freshly collected eggs in 30 mL FSW. On Days 1 and 2, only ∼100 eggs were available for exposure to sperm. On the other days, a surplus of eggs were collected for exposure.

On each subsequent day, we mixed the same amount of stored sperm with as many eggs as we could collect and estimated the total number of fertilized and unfertilized embryos. We found that 2 × 10^7^ sperm consistently fertilized ∼150 eggs after 3, 5, and 6 d at 4°C. On days 1 and 2, we were only able to collect ∼95 eggs, nearly all of which were fertilized. These latter data were consistent with the notion that 2 × 10^7^ sperm could fertilize ∼150 eggs. In these experiments, all embryos developed and metamorphosed into normal juvenile colonies. These results suggest that it is possible to store sperm for at least 6d at 4°C without an appreciable drop in fertilization capability, thereby enabling shipment of sperm samples. This also demonstrated that sperm motility is not necessarily a good predictor of fertilization success when gametes are mixed under controlled conditions.

### Determining cryoprotectant toxicity to sperm

We tested the acute toxicity of three common cryoprotectants (DMSO, methanol, and glycerol). Sperm incubated with the three concentrations (5, 10 and 15%) of DMSO or methanol displayed comparable motility after 30 min. In contrast, sperm exposed to 15% glycerol ceased moving immediately, while those exposed to 10% and 5% glycerol were non-motile within 30 min.

To determine whether cryoprotectant-treated sperm would be able to fertilize eggs, we exposed sperm to each cryoprotectant for 30 min and mixed 4.1 × 10^6^ sperm with 40 freshly collected eggs in a total volume of 50 mL. Sperm exposed to 10% or 15% of any cryoprotectant were unable to fertilize eggs. In contrast, sperm treated with 5% of any cryoprotectant yielded 3-12 embryos (Fig 5). From this, we concluded that 5% DMSO or methanol would be suitable cryoprotectants.

**Fig 5.**
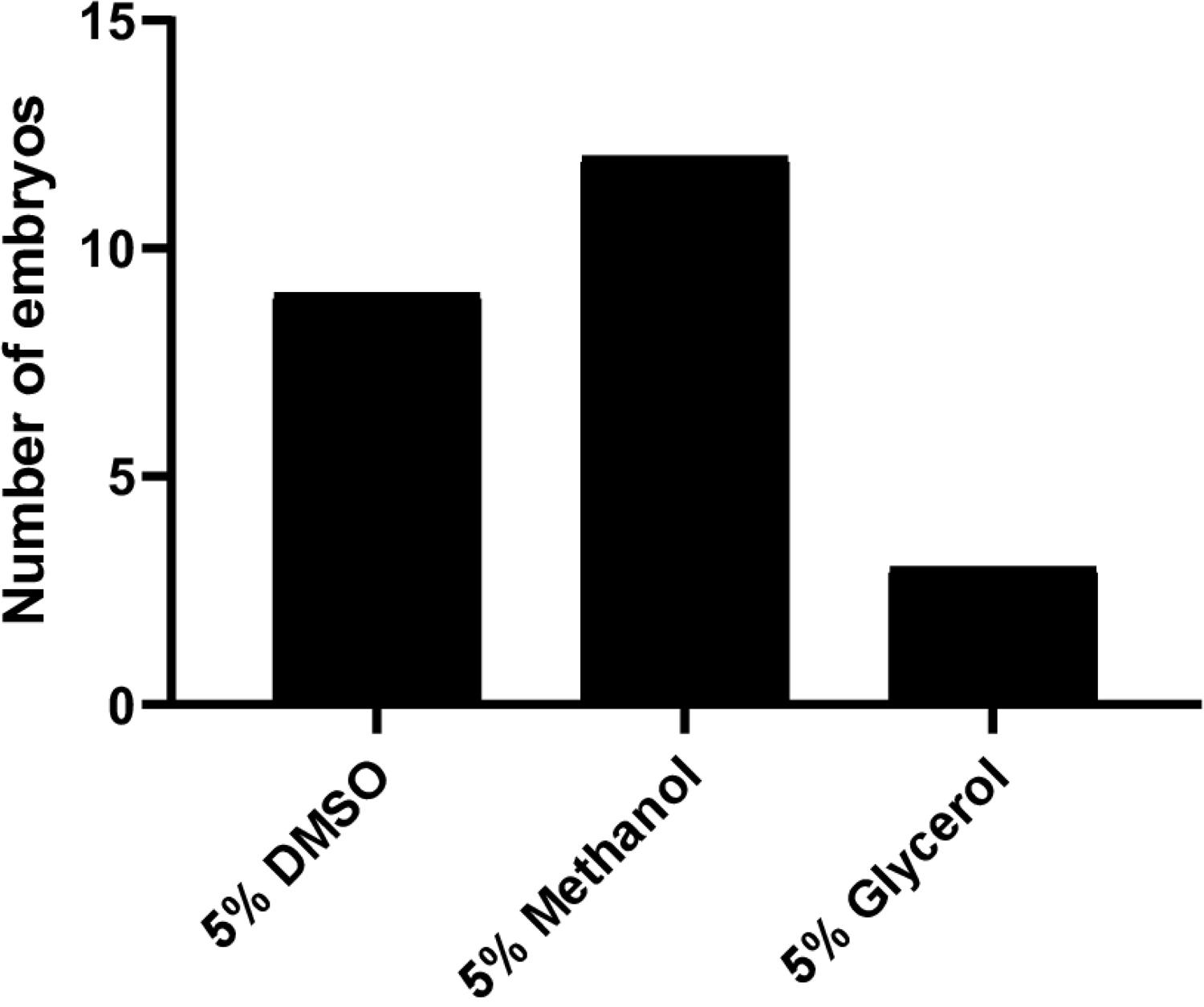
Number of fertilized eggs using cryoprotectant-treated sperm. For each condition, 30-40 eggs were exposed to 4.1 × 10^6^ sperm in a total volume of 50 mL.

### Identifying suitable freezing conditions

While many factors affect the quality of cryopreserved sperm, three key parameters must be balanced: cryoprotectant concentration, sample concentration, and cooling rate. For example, higher cryoprotectant concentrations can be more toxic, whereas lower concentrations may not sufficiently protect the cells. Moreover, the toxicity of a given concentration of cryoprotectant often decreases as the sample concentration increases [10]. The cooling rate must also be slow enough to allow cells to dehydrate sufficiently (to minimize intracellular ice formation), but fast enough to freeze them before concentrations of intracellular salts or pH (i.e., solution effects) or the cryoprotectant become damaging.

To survey the effects of freezing rate on sperm in either 5% DMSO or 5% methanol, we cooled sperm at 5°C/min and 30°C/min (Table 1, Experiment 1; Fig S2, S3). Samples were stored in liquid nitrogen for at least 21 hr before they were thawed and evaluated. In all conditions, the concentration of intact sperm in the thawed samples was reduced from 1 × 10^7^ to 2 × 10^6^ or fewer, nearly ten-fold, likely due to cell rupture either during freezing or thawing. Overall, between 5 × 10^4^ and 1 × 10^5^ fewer sperm were detected in the 30°C/min samples than in the 5°C/min samples suggesting that the faster rate did not allow sufficient osmotic egress and intracellular ice was formed. We incubated aliquots of each thawed sample with 75 freshly collected eggs. Despite the low numbers of sperm used (≤5 × 10^5^), at least one egg was fertilized in each condition. This indicated the presence of viable sperm and suggested that increasing the effective sperm concentration would increase fertilization.

**Table 1.** Overview of frozen samples and fertilization potential.

|  | Experiment 1 |  |  |  | Experiment 2 |
| --- | --- | --- | --- | --- | --- |
| Cryoprotectant | 5% DMSO | 5% DMSO | 5% Methanol | 5% Methanol | 5% DMSO |
| Initial sperm concentration (sperm/mL) | $1 \times 10^7$ | $1 \times 10^7$ | $1 \times 10^7$ | $1 \times 10^7$ | $5 \times 10^7$ |
| Cooling rate (°C/min) | 5 | 30 | 5 | 30 | 20 |
| Hours stored frozen | 69 | 69 | 69 | 69 | 21 |
| Thawed sperm concentration (sperm/mL) | $2 \times 10^6$ | $1.5 \times 10^6$ | $2 \times 10^6$ | $1 \times 10^6$ | $5 \times 10^7$ |
| **Total sperm mixed with 75 eggs** | $5 \times 10^5$ | $3.8 \times 10^5$ | $5 \times 10^5$ | $2.5 \times 10^5$ | $1.2 \times 10^7$ |
| **Number of embryos** | 2 | 2 | 2 | 1 | 10 |

We increased the volume and concentration of sperm collected by fabricating a sperm collection chamber by 3-D printing that allowed incubation of as many as ten slides bearing male colonies in <100 mL of water, thus eliminating the need to collect sperm with pipettes. This enabled collection of 10^9^ sperm per day (a 100-fold increase). We froze the sperm at a concentration of 5 × 10^7^/mL at a cooling rate of 20°C/min. When thawed, these sperm samples remained at a concentration of 5 × 10^7^ sperm/mL (Table 1, Experiment 2; Fig S4). Moreover, the number of fertilized eggs increased to 10.

These results encouraged us to test whether we could further increase fertilization by increasing the concentration of sperm samples. We froze sperm at five different concentrations ranging from 10^7^ to 10^9^/mL (Fig 6) at the 20°C/min cooling rate (Fig S5) and stored them in liquid nitrogen for 21 hr. The concentration of each sample post-thaw had the same count as before freezing. When thawed, sperm frozen at 10^9^/mL were able to fertilize 270-275 eggs. Sperm (in descending order of concentration) at 5 × 10^8^/mL fertilized 235-250 eggs; 1 × 10^8^/mL fertilized 150-250 eggs; 5 × 10^7^/mL fertilized 26-150 eggs, and 1 × 10^7^/mL fertilized 5-100 eggs. All embryos developed into larvae and were able to metamorphose into a primary polyp with no visual abnormalities. Thus, cooling sperm at a rate of 20°C/min and at concentrations in excess of 1 × 10^7^ showed best fertilization.

**Fig 6.**
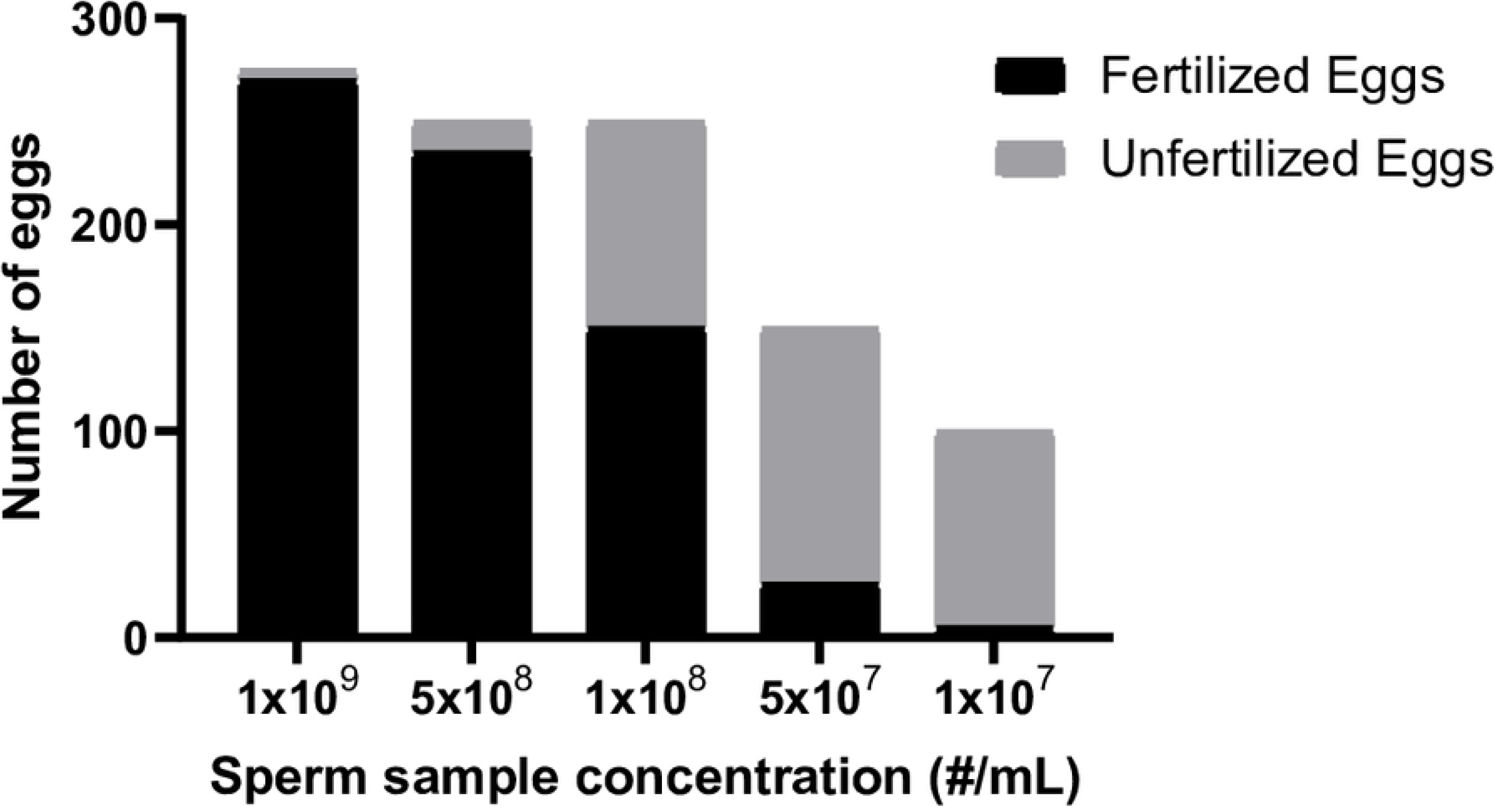
Fertilization comparing frozen sperm. Each thawed sperm sample was exposed to different number of eggs. In each case, the number of eggs collected was manually estimated to be a surplus of what each respective sperm sample could fertilize based on their concentration.

## Discussion

### Sperm motility and viability

The sperm motility and concentration-related phenomena reported herein provide some insight of the basic characteristics of sperm clouds that have not been previously observed. While the results from this feasibility study are promising, there are several improvements and future experiments that can be pursued. There was a large standard deviation (<50%) in motility and concentration between individually collected sperm streams. Part of this variation reflects the imprecise manner traditionally used for collecting sperm as it is released. These findings reinforce the need to standardize collection methods and sperm concentrations. Future studies can also address other outstanding questions related to these characteristics. For example, when and how are sperm activated? Does the sperm concentration affect activation and motility? Determination of how these features could affect cryopreservation, especially among different genotypes, would be useful in expanding and making protocols more robust for *Hydractinia* and potentially other cnidarian models.

Concepts such as these have been studied quantitatively in aquatic species previously at the commercial scale, for example in blue catfish, *Ictalurus furcatus*, (for hatchery production of hybrids) [14] by use of industrial engineering and simulation modeling approaches [15]. Those studies were based on use of automated high-throughput processing [16] developed using commercial dairy industry approaches [17] but are also relevant for processing at lower throughput. The emphasis in such approaches is on the application level for repository development, rather than on the research level for optimization of individual components (e.g., cooling rate or cryoprotectant choice) for protocol development.

Another application-level concept often overlooked in traditional research approaches is the refrigerated storage of samples prior to freezing or use. Such storage enables shipping of germplasm for processing elsewhere, and can avoid waste by identifying the usable working lifetime of valuable material. We tested the retention of *Hydractinia* sperm fertility after storage in FSW at 4°C and found that freshly collected sperm and sperm stored for 6 d could fertilize comparable numbers of eggs. This result suggested that sperm could be stored in FSW at 4°C even longer and still produce viable embryos. Identifying these basic storage conditions is useful in cases when resources are not available to process on-site and samples must be transported to another facility for processing and storage.

Future studies should compare fertility across a range of storage temperatures with longer storage times when appropriate, and couple that with freezing experiments to evaluate the effects of storage on cryopreservation survival. In addition, extender solutions (e.g. buffers) can influence the quality and retention of fertility of sperm during storage [18–20]. Future studies should also address the total fertilization window for eggs. While mixing gametes ≤30 mins post-release has been the community guideline for producing quality embryos, it has not been determined quantitatively and it is possible that storage at a cooler temperature may extend fertility.

### Determining cryoprotectant toxicity to sperm

The acute toxicity assay we performed was at a small scale, but yielded useful information regarding potential cryoprotectants. We initially observed limited fertilization using the treated sperm which demonstrated feasibility and a basis for improvement. In future studies, the potential effects of cryoprotectant toxicity on sperm and egg should be evaluated more clearly. If toxicity is affecting fertilization, sperm can be rinsed to reduce or eliminate the cryoprotectant before exposing them to the eggs. The limited fertilization we observed also emphasized the need to process sperm in concentrations that were relevant to those used for breeding. This prompted the design of a custom 3-D printed collection chamber to improve sperm collection, and enabled evaluation of cryopreservation conditions that resulted in effective post-thaw fertilization rates. This improved collection method provides expanded opportunities for standardized evaluation of cryoprotectants and concentrations, while bearing in mind that such choices should be governed by overall utility at the process level rather than optimizing singular factors (e.g., motility) at the individual step level. For example, a certain cryoprotectant may yield a slightly lower motility value than other chemicals, but is cheaper, less toxic to sperm cells, and allows more flexibility in timing and cooling rates. In research-driven studies, the highest motility would be recommended; in application-driven work, the cryoprotectant that increases efficiency and reliability would be recommended.

Other benefits of placing a focus on application include that work in the present study can be directly scaled up for use with hundreds of animals and multiple laboratories. Work addressing repository development in previous studies, with blue catfish for example, can be generalized to *Hydractinia* because the approaches used are the same, including the use of French straws that can be filled, sealed, and labelled using automated equipment (e.g., the Minitube Quattro system at the AGGRC can process 15,000 straws per hour). In addition, cryopreservation in *Hydractinia* can be directly transferred from a central facility (such as the AGGRC) to on-site work within an existing laboratory by use of high-throughput mobile cryopreservation capabilities [21], or by establishment of full high-throughput cryopreservation capabilities such as in creation of a central *Hydractinia* Stock Center (for economic analysis, see [22]). Development of in-house cryopreservation capabilities within research laboratories will be greatly strengthened by the recent developments in 3-D printing described above (e.g., [23]) including fabrication of probes for monitoring and storing temperature information [24], and the potential for sharing of open-source design files for production of inexpensive, reproducible freezing devices that can be integrated with strong quality control programs (e.g., [25,26]).

### Identifying suitable freezing conditions

While there are no other *Hydractinia* cryopreservation protocols to directly compare our results to, there are protocols that have been developed for sperm from various coral species which can serve as an indirect comparison for some of the key parameters. One protocol in particular has been instrumental in banking the germplasm of 31 coral species from around the world [6,27,28] and additional protocols have been developed in two coral species [29,30]. Briefly, we can compare our method with the cryoprotectant, container, and cooling methods of these studies. Similar to two of the studies [28,30], DMSO was used as the cryoprotectant but at higher final concentrations (≥10%), and in the other [29] 20% methanol was used with 0.9 M sucrose as an extender. These cryoprotectant concentrations are higher than what our trials suggested would be suitable for *Hydractinia* sperm, however there are two major differences that may explain this discrepancy and proffer improvements to this study. The *in vitro* fertilization in this study used a volumetric sperm to egg ratio of 50 mL with ∼10^5^/mL sperm to 30-40 eggs, which was considerably more dilute in comparison to each coral study in which the ratios used to determine post-cryoprotectant fertility were 5 mL with 10^6^/mL sperm to 30-50 eggs [28], 1 mL with 10^5^/mL sperm to 20 eggs [29], or 4 mL with 1.5 × 10^7^/mL sperm to 50 eggs [30]. In future acute toxicity assays, optimizing the volumetric sperm to egg ratio (in our case, reducing the volume and increasing the sperm concentration) would improve the assessment of acute toxicity before moving onto freezing. Previous studies with eastern oyster *Crassostrea gigas* have shown that much of the variation in sperm cryopreservation response is procedural rather than biological (e.g., “male-to-male variation”), and control of sperm concentration is necessary for reproducible results [31].

One of the coral protocols cryopreserved 1-mL samples in 2-mL cryovials [28], whereas the other two studies [29,30] cryopreserved samples in 0.25-mL French straws. French straws offer several advantages over traditional cryovials. French straws require less storage space and can be easily processed manually in the case of a few samples, or more efficiently in high-throughput with automated filling, labeling, and sealing for hundreds to thousands of samples. In addition, samples can generally be cooled in French straws at a faster rate than in cryovials, in large part due to the higher surface-area-to-volume ratio of straws (which can also decrease variability during freezing). In cryovials, there is potentially more variation across the sample volume as material on the periphery could freeze more rapidly than that closer to the center. Also, vials typically have thicker walls with greater insulative potential slowing heat removal from the sample.

With regard to cooling rates, there are several differences which make these studies difficult to compare. First, the equilibration temperature and time used were slightly different [28,30] or not explicitly quantified [29]. Second, the ending temperatures used to calculate the freezing curve were different where one study used -80°C [28] but the other two used the coldest achievable temperature between -110°C and -130°C. Theoretically, the ending temperature should not affect the rate calculation if the freezing rate is constant, but unless the temperature is monitored while the samples are being frozen, fluctuations are difficult to account for. Although the different procedures make studies difficult to compare, it is critical that all details surrounding the freezing process be documented for quality samples and reproducible results [11]. For this reason, only two of the studies can be referenced for reproducibility and generally compared in relation to their cooling rate [28,30]. Both studies used an equilibration temperature between 24-29°C and equilibration time of 15 [30] or 20 min [28] where in this study the equilibration temperature was 4°C for 20 min. The selection of 4°C as the equilibration temperature in our study was in part due to the usage of a controlled-rate freezer.

One obvious difference among the present study and the three published coral studies is that each used suspension at defined heights above liquid nitrogen to freeze samples. This method is difficult to standardize, and is less precise than using a controlled-rate freezer. However, it has significant advantages including affordability, availability, and portability. In future studies, a comparison of samples frozen at comparable nominal rates by various methods should be done to enable harmonization of results and reporting, providing multiple options for freezing that could be selected based on the needs of the user. Other factors that could be investigated in future work include whether offspring produced from cryopreserved sperm will mature into full adults and whether the male-to-female ratio is affected.

### Approaches to repository development for aquatic species

Recent advances in consumer-level technology provide opportunities to custom-design open-source options for hardware and other tools necessary to assist repository development beyond that provided by adaptation of traditional livestock practices. Customizing the design of the 3-D printed collection chamber greatly increased the efficiency and success in identifying suitable freezing conditions. To collect a useful number of sperm, the standard collection method via Pasteur pipette or micropipette is labor intensive, and poses logistical problems in the case of multiple collectors. Given our previous approach, collecting all sperm would be possible but would require filtering all the water from the bin (∼2 L) or having access to a large capacity centrifuge. Thus, by customizing a chamber to minimize the collection volume (<100 mL) and maximize the total yield of sperm (as many as ten slides bearing *Hydractinia*), we were able to directly improve and standardize processing efficiency.

In addition, custom design of devices is also possible for freezing activities. The polylactic acid (PLA) used for 3-D printing does not become brittle or stiff as do other plastics when exposed to cryogenic temperatures (such as liquid nitrogen) [32] making 3-D printed objects safe and useful for such applications. Various devices can typically be fabricated at low cost (e.g., $10 or less for material costs) using consumer-level printers ($250 or less) that offer high resolution, flexibility, and short learning curves. There are large internet-driven user communities for these printers and thousands of videos (such as on YouTube) are available for printer set up, training, and troubleshooting. In addition, design files can be shared on a number of sites (e.g., Thingiverse, and Github) for others to print and customize. In this way, devices used in cryopreservation and repository development can be developed, shared, and standardized within research communities, greatly reducing costs of cryopreservation, and making reliable methods widely available. Systems such as this must be accompanied by quality control and quality assurance programs, however, to ensure that samples meet minimum thresholds for repository use [25,26].)

Overall, the success in the present study of using a generalizable approach for *Hydractinia* sperm provides further evidence that cryopreservation protocols are not necessarily species-specific. For example, a single generalized protocol was applied to more than 20 species within the genus *Xiphophorus* and two other species in the genus *Poecilia* to enable repository development to safeguard the genetic resources of these valuable biomedical model species [12]. Overall, more research is needed for aquatic species in general to quantitatively assess factors important to practical repository operation with cryopreserved sperm (e.g., [14]), and standardization of procedures and reporting is necessary to enable meaningful comparisons across studies [11]. The present study offers evidence that substantial repository-level benefits can be realized by generalizing cryopreservation at the application level, rather than trying to optimize new protocols on a species-by-species basis, and restricting this work to the traditional (reductionist) research level [11].

## Conclusions

This feasibility study showed that it is possible and a worthwhile endeavor to pursue *Hydractinia* sperm cryopreservation as a long-term storage option for genetic resources. Specifically, we demonstrated that sperm cooled at 20°C per min in 5% DMSO at a concentration of 10^8^-10^9^/ml in 0.25-mL French straws were able to fertilize 150-300 eggs, which developed into juvenile colonies. In our experience, a population of 150 juvenile colonies typically contains sufficient numbers to establish a strain for propagation via asexual reproduction (i.e., they will grow into healthy adults with the genotype of interest) or breeding to produce subsequent generations. With some additional work it should be possible to reliably freeze and re-derive specific genotypes. This would greatly enhance the utility of *Hydractinia* as a model system for cnidarian genetics. In addition, we expect this work could also provide a guide to researchers seeking to develop cryopreservation approaches in other cnidarian species.

While this study has direct implications for the *Hydractinia* community, there are several considerations that can be discussed with regard to communities that work on other cnidarian models. Lack of long-term storage options has been one of the limitations to nearly all cnidarian research. Cryopreservation has not been pursued either due to the lack of resources to achieve and maintain frozen samples, or the lack of necessity as many cnidarian models can be cultured relatively simply and the animals can regenerate. A notable exception to this are cryopreservation efforts for conservation in coral species and their symbionts due to importance of corals to reef biodiversity, and the overall decline in health and prevalence of corals globally over the past several decades [33–36].

Another emerging problem for popular research models is the rapid proliferation of new lines and mutants that would require maintenance as live animals which is expensive and risky without cryopreservation. With these common limitations in mind, cnidarian communities need to come together and agree on a consistent and foundational approach towards cryopreservation of all cnidarian models for the ultimate purpose of repository development and establishment of repository networks. By having this long-term goal in mind, we can more systematically work towards developing, protecting, maintaining, distributing, and utilizing an expanding pool of cnidarian genetic resources.

A centralized repository or stock center is a necessity for well-developed research organisms and part of the success with these has been due to collaboration among laboratories and the sharing of tools, systems, and resources throughout the communities. For example, mouse resources are largely centralized with The Jackson Laboratory (https://www.jax.org/), zebrafish databases and lines are found in the Zebrafish International Resources Center (ZIRC, University of Oregon), *Drosophila* utilizes the Bloomington *Drosophila* Stock Center (BDSC, Indiana University Bloomington), *Caenorhabditis elegans* and other worm-related models localize their resources in WormBase (wormbase.org/), and *Xenopus* related resources are found in Xenbase (https://www.xenbase.org). Having a wealth of such resources and information available to these communities makes these model systems much more useful and available to investigators, whereas model systems that require development of basic tools can be more challenging on many levels.

Future studies should establish a standardized approach for the storage, shipment, and use of frozen *Hydractinia* samples that can be made available throughout the research community. Current models for this would include development of repositories or a repository system, and the potential incorporation of these entities into a community-based stock center. An existing model for such organization exists in ZIRC which maintains more than 43,000 research lines of zebrafish as frozen sperm (https://zebrafish.org/). In addition, to assist standardization of protocols and approaches, it may be useful to establish community-level mechanisms to design and share inexpensive devices that can be used to support users across a wide range of experience and skill levels in culture, spawning, and cryopreservation of *Hydractinia*. Lastly, cryopreservation and repository development should be expanded to include additional germplasm and somatic cell types.

## Acknowledgements

We thank Mallory Lemoine, William Childress, Amy Guitreau, Liu Yue, Teresa Gutierrez-Wing of the AGGRC for technical assistance and discussion.

## Supporting Information Captions

**Figure S1. Pedigree of the colonies used to generate germplasm and offspring**. Field-collected colonies are denoted with black symbols. Colony 291-10 is the offspring of two colonies collected from Lighthouse Point, New Haven, CT in 2014. Colony 295-8 is the offspring of a field collected colony and a laboratory strain, 235-33. The pedigree of colony 235-33 can be recreated by concatenating previously published pedigrees (shaded area) (Cadavid et al. 2004; Powell et al. 2007). Colony AP100-88 is from the mapping population in Powell et al. (2007). Colony 431-44 is from the mapping population in Cadavid et al. (2004).

**Figure S2. Experiment 1 cooling curve (5°C/min)**.

The first trial included a slow cooling rate of 5°C/min with a sperm concentration of 1×10^7^/mL. The green line represents the programmed temperature profile, red is the chamber temperature, and blue is the sample temperature. Samples were held at -80°C for at least 5 min before plunging into liquid nitrogen.

**Figure S3. Experiment 1 cooling curve (30°C/min)**.

The first trial included a fast freezing rate of 30°C/min with a sperm concentration of 1×10^7^/mL. The green line represents the programmed temperature profile, red is the chamber temperature, and blue is the sample temperature. Samples were held at -80°C for at least 5 minutes before plunging into liquid nitrogen.

**Figure S4. Experiment 2 cooling curve (20°C/min)**. The second trial adjusted the freezing rate to a moderate rate of 20°C/min with a more concentrated sperm sample (5×10^7^/mL). The green line represents the programmed temperature profile, red is the chamber temperature, and blue is the sample temperature. Samples were held at -80°C for at least 5 minutes before plunging into liquid nitrogen.

**Figure S5. Cooling curve of various sperm concentrations at (20°C/min)**. Each sperm concentration (10^7^-10^9^/mL) was frozen at the same time at 20°C/min. The green line represents the programmed temperature profile, red is the chamber temperature, and blue is the sample temperature. Samples were held at -80°C for at least 5 min before plunging into liquid nitrogen.

**Table S1. Slicer software settings used to 3-D print collection chamber**.

**Table S2. Printer hardware features**.

